# LAVR-289, a New Orally Bioavailable Inhibitor of Adenovirus Replication *in vitro* and *in vivo*

**DOI:** 10.1101/2025.02.20.639236

**Authors:** Charlotte Quentin-Froignant, Ann Tollefson, Anna Cline-Smith, Sandrine Kappler-Gratias, Nicolas Landrein, Vincent Roy, Luigi A. Agrofoglio, Karoly Toth, Harald Wodrich, Franck Gallardo

## Abstract

Adenoviruses are responsible for a range of pathologies, including respiratory infections in children, accounting for 5-10% of such cases. Although most adenovirus infections are self-resolving, they can cause serious illness, particularly in immunocompromised individuals. There is currently no approved treatment for adenovirus infections, although various therapeutic approaches are under investigation, including nucleoside analog inhibitors of replication. However, these treatments have shown limited efficacy. In this study, we report on the antiviral activity of LAVR-289, a broad-spectrum acyclonucleoside phosphonate exhibiting potent *in vitro* efficacy against several adenovirus serotypes, comparable to that of brincidofovir. LAVR-289 specifically inhibits viral replication, blocking the formation of viral replication centers and preventing late protein expression without affecting viral entry or delivery of viral genomes to the nucleus. *In vivo* using immunocompromised Syrian hamsters infected with HAdV-C6, oral administration of LAVR-289 resulted in 100% animal survival. These results suggest that LAVR-289 holds promise as a potential therapy for adenovirus infections, particularly in immunocompromised patients.

**Highlights:**

- LAVR-289 is a unique acyclic nucleoside phosphonate prodrug.
- LAVR-289 displays antiviral activity against Adenovirus with an EC_50_ of about 100 nM against HAdV-C5 ANCHOR.
- LAVR-289 inhibits viral replication by targeting viral DNA polymerase, preventing DBP clustering and replication center formation.
- In immunosuppressed Syrian hamsters, LAVR-289 is well tolerated and ensures 100% survival while effectively stopping virus replication.
- LAVR-289's broad-spectrum activity positions it as a promising treatment for immunocompromised patients facing multiple viral infections.

## Introduction

Adenoviruses are small (∼90nm), non-enveloped DNA viruses of the *Adenoviridae* family^1^. Their distinctive icosahedral structure allows them to remain stable in the environment while retaining their infectivity. Adenoviruses infect a broad range of hosts including humans and animals. In humans, more than 100 types have been identified and classified into seven species (from A to G). They induce a broad variety of pathologies reflecting the cellular tropism of the different species^2^. Adenovirus infection is primarily known to cause respiratory infections in children causing 5 to 10% of pediatric respiratory infections, while being responsible for almost 20% of pneumonia cases in newborns and 1-7% of cases in adults^3^. Adenovirus infections can also cause other pathologies, such as gastroenteritis and keratoconjunctivitis. Fatal systemic infections are mainly seen in immunocompromised persons, albeit they have also been reported for immunocompetent patients^4,5^ .

Although in general, adenovirus infections are self-resolving, prevention and treatment of adenovirus infection, especially for immunosuppressed people, remains a challenge. An efficacious vaccine has been developed, but its use is limited to military personnel applications^6^. To date, there is no FDA or EMA approved treatment for adenovirus infection. Different therapeutic solutions have been investigated over the past years, such as natural products, epigenetic modulators and steroid-based compounds for example^7–9^. Nucleoside/nucleotide analogs show remarkable *in vitro* activities and possess a strong barrier to resistance by targeting the viral polymerase. One such compound, cidofovir (CDV), an acyclic nucleoside phosphonate, has been administered off label in the clinic for decades^10,11^. However, CDV suffers from poor cellular uptake and nephrotoxicity. Its lipidic prodrug brincidofovir (BCDV) represents a major advance in therapy against adenovirus and has been successfully used to control adenovirus viremia in hematopoietic cell transplant patients^12^. Unfortunately, oral administration of BCDV caused dose-limiting gastrointestinal toxicity and the compound failed a Phase III clinical trial^13^.

We have recently developed LAVR-289, a [(*Z*)-3-(acetoxymethyl)-4-(2,4-diaminopyrimidin-6-yl)oxy-but-2-enyl]phosphonic acid prodrug with broad-spectrum anti-DNA virus activity^14^ and reported a formulation for the compound^15^. LAVR-289 can inhibit *in vitro* the activity of varicella-zoster virus, human cytomegalovirus, vaccinia virus, and human herpesvirus 1. In this study, we show that LAVR-289 displays nanomolar activities against several HAdV serotypes, being at least as effective as brincidofovir *in vitro*. LAVR-289 specifically impacts viral DNA replication, resulting in the absence of late protein expression while early events of virus infection are not affected. *In vivo* administration *per os* once daily in Syrian Hamster infected with a lethal challenge of HAdV-C6 showed 100% survival, several log reduction of adenovirus replication in the liver and protection of organ function.

## Methods

### Antiviral compounds

LAVR-289 was synthetized at a purity of 98% (purity determined by HPLC area) according to the procedure described previously by our team^14^. A 10mM stock solution was prepared in DMSO and stored at -20 °C before use. BCDV was ordered from Clinisciences (100mg, ref 444805-28-1).

### Cells and viruses

HEK293 (ATCC CRL-1573), human foreskin fibroblast (HFF) and HFF stably expressing the adenovirus receptor CAR (HFF-CAR, kindly provided by J. Dechanet, Univ. Bordeaux) were used in this study. Cell lines were cultured in Dulbecco’s Modified Eagle’s Medium (DMEM) supplemented with 10 % fetal bovine serum (FBS) (Eurobio-Scientific), 1mM sodium pyruvate (S8636; Sigma Aldrich), L-Glutamine (G7513; Sigma Aldrich) and Penicillin-Streptomycin solution (P0781; Sigma Aldrich). Cells were grown at 37°C with 5% CO2. HAdV-C5 ANCHOR GFP virus^16^, HAdV-C5 (ATCC VR-1516) and HAdV-C6 (Strain Tonsil 99; ATCC VR-6) were used. HAdV-B3 (gift of Vivian Mautner, University of Birmingham), HAdV-B7 (gift of Kathy Molnar-Kimber, Wyeth), HAdV-C1 (ATCC VR-1078), HAdV-C5 (ATCC-VR-5, Adenoid 75), HAdV-C6 (ATCC VR 1083, Tonsil 99), HAdV-D9 (ATCC VR-10, HAdV-E4 (RI-67, CDC), and HAdV-F41 (ATTC VR-930, TaK) were used for the experiment shown in Table 1.

**Table 1:**
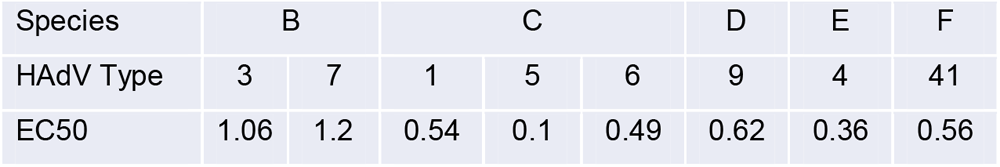
EC_50_ of LAVR-289 for Different HAdV types, μM (HFF Cells)

### High content imaging and quantification

High content imaging and quantification was described previously^17^. Cells were infected with HAdV-C5 ANCHOR at the indicated MOI and fixed at the indicated time post infection for 15 min with formalin. Cells were washed in PBS and stained with Hoechst 33342 (1µg/mL) for 10 min and processed for high content microscopy. We used a Thermo CellInsight CX7 microscope with the indicated objective for image acquisition. Compartmental analysis was used to calculate the infection rate (number of ANCHOR positive cells over the total number of cells) and replication level (integrated intensity of the replication centers in infected cells). Combenefit was used to calculate EC50s of the different tested products^18^.

### HAdV serotype testing

HFF cells were plated at 10^4^ cells/well on 96-well plates one day before infection. On the day of infection, LAVR-289 was serially diluted in a 1:3 series on dilution plates as paired replicates and added to HFF wells on the cell plates. The highest final concentration of LAVR-289 was 20µM in the first row on the cell plates. Viruses were diluted appropriately for each serotype. Viruses were added immediately subsequent to drug additions. Cells were fixed in 3.7% paraformaldehyde in PBS at 2 days post-infection (HAdV-C1, -C5, -C6, -B3, -B7, and -E4) or at 3 days post-infection (HAdV-D9 and HAdV-F41). Cells were subsequently stained with anti-hexon antibody (mouse monoclonal 2HX-2) and secondary anti-mouse IgG-HRP conjugated antibody, visualized by substrate TrueBlue, and quantified on a CTL Elispot reader. EC_50_ values were calculated in GraphPad Prism.

### Infection and immunofluorescence

HFF-CAR cells were grown on coverslips in a 12-well plate at a density of 2×10^5^ cells on the day of infection. A stock solution of 10mM LAVR-289 in DMSO was diluted in DMEM and cells were pre-incubated with compound at the indicated final concentrations or the respective vehicle control for 1 hour prior to infection. Infections were carried out for 30 min using ∼3000 physical particles of HAdV-C5 per cell (for the 3h time point) or using a 1:1 mixture of HAdV-C5 and E1-deleted HAdV-C5-ANCHOR (for the 24h time point) at the same concentration before the inoculum was removed. Cells were incubated for 3h or 24h in 1ml medium in the presence of LAVR-289 at the indicated final concentration. Coverslips were subsequently fixed using 4% paraformaldehyde (pH□=□7.4) and processed for immunofluorescence (IF) or alternatively cell pellets were collected for western blot (WB) analysis. For IF analysis, cells were blocked/permeabilized using IF buffer (PBS+10, fetal calf serum (FCS) +0.1% saponin). Primary and secondary antibodies were diluted in IF buffer and applied to cells for 1 h in a humid chamber at 37°C. Coverslips were mounted using fluorescence mounting medium (Dako) containing 1□μg/ml DAPI (Sigma) to counterstain nuclei. Cells were imaged using a confocal microscope (Leica) at 200 ms for each channel using a z-series of 3 planes a 0.3 µm and analyzed using ImageJ software. For genome quantification, we generated z-projections of 8-11 independent fields of view amounting to 57-83 cells per condition. Genomes were quantified using imageJ macros to determine the number of genomes per nucleus^19^.

### Antibodies

To detect delivered virions, cells were stained with mouse anti-protein VII monoclonal antibodies at 1:100 dilution (produced in the lab^20^) and mouse anti-DBP antibodies at a dilution of 1:100 were used to detect DBP and replication centers. Secondary antibodies from donkey coupled with respective Alexa Fluor were used at a dilution of 1:500 (Life Technologies). For the WB analysis we used the same primary anti-DBP antibody diluted 1:500 as well as rabbit anti-HAdV-C5 serum at a 1:5000 dilution (both kindly provided by R. Iggo, Univ. Bordeaux) and peroxidase-coupled secondary antibodies (Jackson Immuno Research) diluted 1:10,000.

### Drug formulation for in vivo experiments

LAVR-289 was obtained from the National Institutes of Allergy and Infectious Diseases (Bethesda, MD, USA) in semi-liquid form and stored at -20°C. For *in vivo* administration, it was dissolved in 10% DMSO, 10% Cremophor, 80% PBS at a concentration that contained the required dose in 1 ml volume. The drug was made up once, aliquoted into daily portions, and stored at 4°C. Aliquots were allowed to equilibrate to room temperature before dosing.

### In vivo toxicity

Approximately 100 g male hamsters were purchased from Envigo and immunosuppressed using intraperitoneal administration of cyclophosphamide (CP) at a single dose of 140 mg/kg followed by twice weekly injections at a dose of 100 mg/kg. To determine the Maximum Tolerated Dose (MTD), 6 hamsters for each of the 25, 50, and 75 mg/kg dose levels were dosed orally (p.o.) once a day (q.d.) with 1 ml volume of the appropriate solution and were observed and weighed daily. The animals were sacrificed after 15 days of drug administration and gross pathologic observations were made. Serum samples were collected for analysis of plasmatic parameters such as transaminase, creatinine, and blood urea nitrogen levels.

### In vivo efficacy

Approximately 100 g male hamsters were purchased from Envigo. The animals were distributed into 7 groups of 15 hamsters each, with the exception of Group 2 (LAVR-289 50mg/kg alone - uninfected), which consisted of 9 animals. All hamsters were immunosuppressed using cyclophosphamide (CP) intraperitoneally at an initial dose of 140 mg/kg, and then twice weekly at a dose of 100 mg/kg. The hamsters were injected i.v. with vehicle or 2×10^10^ plaque-forming units (PFU)/kg of HAdV-C6 (Lot# 190729), and then treated with drug vehicle or LAVR-289 *per os* (p.o. q.d.) at 5, 17 and 50mg/kg dose levels. A control group received cidofovir (a 37 mg/kg initial dose, and then 20 mg/kg, 3 times weekly). For all groups, drug administration started one day before challenge and continued for the duration of the study. The body weights and any signs of morbidity of the animals were recorded daily. At 5 days post challenge, 5 hamsters (designated at the start of the experiment) of each group except Group 2 were sacrificed, and gross pathological observation was performed. Serum and liver samples were collected to determine transaminase levels and virus burden, respectively. The remaining 10 hamsters were sacrificed at 14 days post challenge. Hamsters that became moribund before Day 14 were sacrificed. Besides animals judged moribund by observation, all hamsters that lost more than 20% of their original body weight were sacrificed, triggering the cessation of the weight curve for the group in which an animal died.

## Results

### LAVR-289 inhibits replication of human adenoviruses at nM concentration *in vitro*

LAVR-289 (Fig 1A) is a new acyclonucleoside phosphonate having remarkable antiviral activities on viruses from the poxvirus and herpesvirus family (see Marcheteau et al., Kappler-Gratias et al., same issue). To investigate if this compound can be active on other dsDNA viruses, we tested its efficacy on HAdV-C5 ANCHOR infected cells. HAdV-C5 ANCHOR GFP virus has been previously developed and characterized to allow visualization of infection and replication in living cells^16^. HAdV-C5 ANCHOR virus is based on an E1-deleted viral vector and grows in E1-complementing HEK293 cells. Infecting HEK293 cells with HAdV-C5 ANCHOR virus at MOI 0.2 for 24h allow the visualization of infected cells (GFP positive) and visualization of replication centers, visualized by the accumulation of GFP fluorescent foci in the cell nucleus (Fig. 1A, left/right). By using a compartmental algorithm, we can quantify the infection rate (number of GFP positive cells/total number of cells) and replication rate (surface area and intensity of replication centers) as a surrogate measurement for HAdV replication. At 125nM, LAVR-289 caused a decrease in virus replication center surface area, whereas at 250nM, replication centers are almost undetectable. To see if inhibition is stable over time, as adenoviruses replicate rapidly in cell culture, we repeated this experiment this time measuring at 48h PI and calculated EC50 using a compartmental analysis to quantify the integrated intensity of replication centers inside infected cells (Fig. 1B, left). In this experiment, we can visualize a dose dependent inhibition of fluorescence accumulation, with pronounced inhibition at concentrations above 300 nM (Fig 1B right). Whereas in the absence of the drug, replication centers display a fluorescence signal of around 940000 fluorescence units on average, replication center signal is decreased to 30000 on average for concentrations above 250nM, indicating a 97% decrease in viral replication level with an EC50 of 90 nM. To compare LAVR-289 against the gold standard BCDV, we infected cells at MOI 0.1 with HAdV-C5 ANCHOR virus and scored infection rate and viral DNA replication level 48h PI. As shown in Fig. 1C, both compounds were able to inhibit viral replication and decrease infection rate at similar EC50s, indicating that LAVR-289 is as effective as BCDV *in vitro*. To test if LAVR-289 can be active on HAdV of different serotypes, we calculated its EC50 on human foreskin fibroblasts infected with HAdV belonging to 5 different species and quantified the suppression of virus replication by measuring hexon expression (Table 1). The HAdV hexon protein is expressed at the late stage of replication, after viral genome replication; thus, it is a valid surrogate of virus replication^21^. LAVR-289 was active against all adenoviruses tested, with EC50s ranging from 100 nM to 1.2 µM. LAVR-289 also displayed a 90% reduction in HAdV-F40 infectious virus production as assessed by Tissue Culture Infectious Dose 50 (TCID50 experiment) (*data not shown*). Altogether, these results show that LAVR-289 is a new inhibitor of HAdV replication *in vitro* with EC50 comparable to the gold standard BCDV. Importantly, LAVR-289 inhibits the replication of all HAdV tested so far.

**Figure 1:**
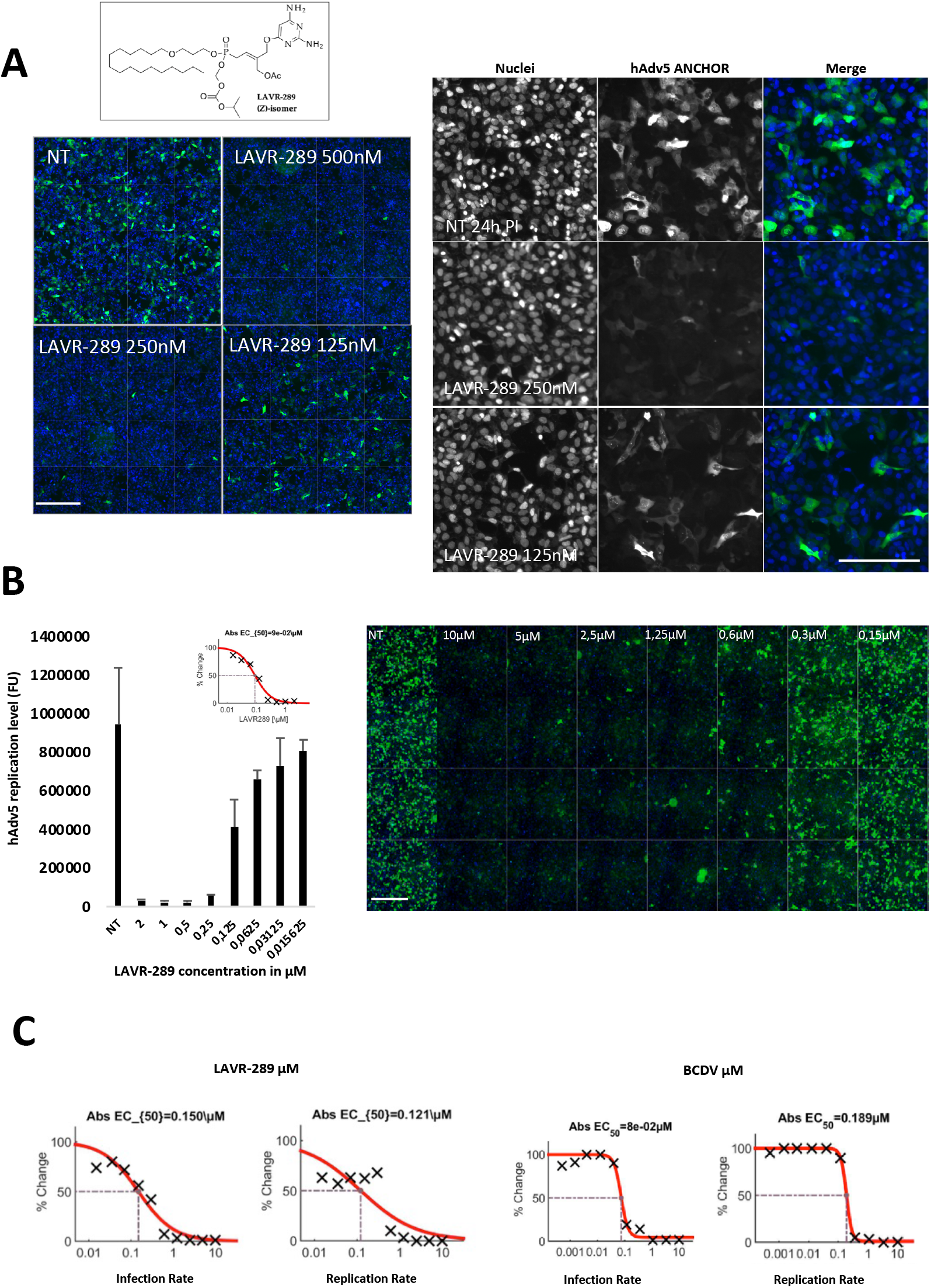
LAVR-289 inhibits replication of human adenovirus 5 at nanomolar concentrations. A) HEK293 cells infected with HAdv5 ANCHOR at MOI 0.2 treated or not (NT) with LAVR-289 at 500, 250 and 125nM concentrations, fixed and imaged 24h PI on a Thermo CX7 CellInsight HCS microscope (10X objective). Merge picture of Hoechst 33342 staining (blue) and Adv5 ANCHOR-GFP (green) are presented. 16 fields reconstruction (left, bar=800µm) and single field (right, bar=400µm) are shown. Note the massive drop in virus replication center surface and intensity when cells are treated with LAVR-289. B) Left, hAdv5 ANCHOR replication level according to LAVR-289 concentrations. Results are presented as mean +SD. Calculated IC50 is 90nM as assessed using Combenefit. Right, single field example of dose response testing of LAVR-289 on HEK293 cells infected with HAdv5 ANCHOR MOI 0,2 at 48h PI. Concentrations of LAVR-289 are indicated (bar=400µm). C) Quantification of LAVR-289 efficiency on the infection rate (left) and replication rate (right) according to MOI at 48hPI. BCDV is used as positive control.

### LAVR-289 inhibits virus propagation by inhibiting early to late replication phase transition

We have shown that LAVR-289 targets the poxvirus viral DNA polymerase (see Marcheteau et al. same issue). To investigate the compound’s mechanism of action against HAdV, we determined its impact on viral nuclear import as an endpoint of the entry phase, replication center formation as a marker for replication onset, and expression of late genes during the post replication phase. For this, we infected HFF-CAR cells with HAdV-C5 in the presence of 200nM and 500nM of LAVR-289 or vehicle alone. At 3 h PI, cells are fixed and processed for immunofluorescence of the viral pVII protein that is bound to incoming viral genomes and which drives viral import into the nucleus^22^. Genome import in the nucleus can be detected via individual pVII foci that correspond to the position of single viral genomes (Fig. 2A). Quantification of the number of pVII foci per nucleus in non-treated (NT) and LAVR-289 treated cells shows that presence of LAVR-289 does not modify HAdV genome import in the nucleus even at the highest concentration of 500nM (Fig. 2B). To assess the effect of LAVR-289 on the onset of virus replication, we co-infected HFF cells with a 1:1 mixture of HAdV-C5 ANCHOR GFP and replicative HAdV-C5 virus (Fig. 2C) at an excess. Co-infection is needed as the HAdV-C5 ANCHOR GFP virus is E1-deleted and needs to be trans-complemented by a WT virus providing E1 to initiate replication. We first checked the formation of replication centers via the expression and clustering of DBP, the viral ssDNA binding protein required for virus replication^23^. In untreated conditions at 24h PI, DBP is expressed and clusters in large replication centers foci, indicating active replication of the virus, confirmed by the accumulation of ANCHOR GFP foci marking replicated viral DNA in the nucleus (Fig. 2C)^16^. In LAVR-289 treated cells, albeit DBP is still expressed to high level, the protein does not cluster and replication centers do not form. Consequently, no replicated viral DNA could be observed in the ANCHOR GFP channel. Moreover, by extracting total proteins of untreated and LAVR-289-treated cells and performing Western-blot using anti-HAdV serum detecting late proteins, we could visualize a strong drop in late protein accumulation, starting at 100nM LAVR-289 (Fig. 2D). As late protein expression requires genome replication, this further indicates that LAVR-289 targets HAdV DNA replication. In contrast, DBP which is expressed from incoming genomes during the early infection phase, accumulates independently of LAVR-289 in the nucleoplasm (Fig. 2D). Taken together, these results show that LAVR-289 does not impact virus entry using genome delivery as endpoint of the entry process, nor does it impact early gene expression as indicated by unwavering DBP expression. In contrast, dose dependent LAVR-289 treatment abolishes viral DNA replication, replication center formation and DBP clustering, resulting in a defect in late protein expression.

**Figure 2:**
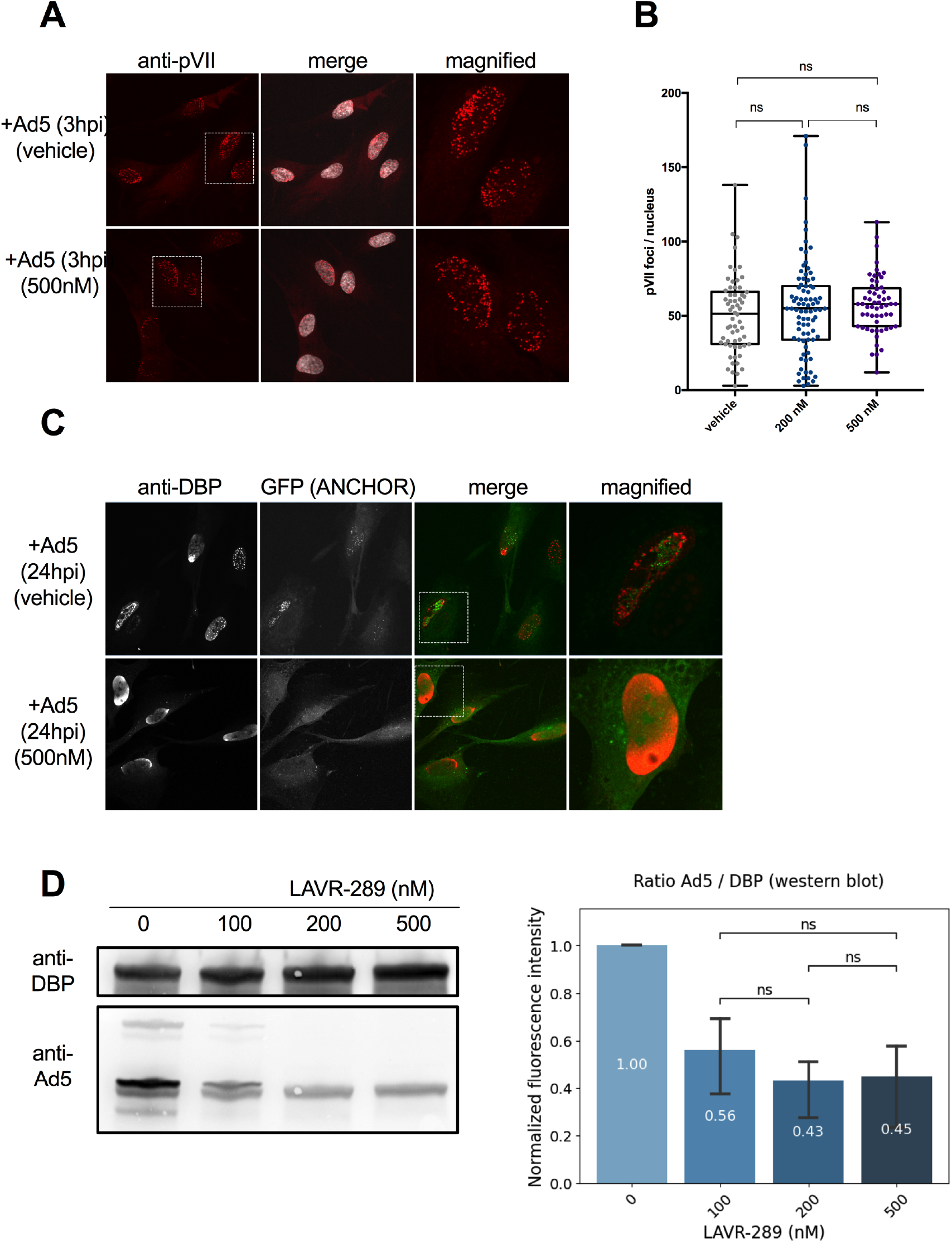
LAVR-289 blocks early replication firing without impacting HAdv5 ANCHOR genome release or early viral gene expression. A) Nuclear mid sections with HAdV-C5 genomes stained with anti-protein VII antibodies at 3 hours post infection in presence of vehicle (top row) or 500nM LAVR-289 (bottom row). Shown is the protein VII stain (first column, red signal) counterstained with DAPI (second column, grey signal) and as magnified image of the boxed area (third column). B) quantifications of nuclear genomes at 3 hours post infection for vehicle control, 200nM and 500nM LAVR-289 concentration (>50 cells per condition, one sided ANOVA, ns = non-significant). C) Nuclear mid sections of HAdV-C5 ANCHOR co-infected cells at 24 hours post infection in presence of vehicle alone (top row) or 500nM LAVR-289 (bottom row). Cells were stained with anti-DBP antibodies detecting replication center (first column, red signal), GFP staining replicated viral genomes (second column, green signal) and as merged signals with (third column) and as magnified image of the boxed area (last column). Note the absence of replication center in presence of LAVR-289. D) Western blot analysis and quantification of HAdV-C5 infected cells at 24 hours post infection treated with LAVR-289 as indicated. Left panel; Equal amounts of lysate were stained with antibodies against DBP expressed prior to replication (top blot) and an anti-adenovirus serum recognizing several structural proteins expressed post replication (bottom blot). Right panel; shows the quantification of the HAdV-C5 total anti-adenovirus signal normalized by the DBP signal (one sided ANOVA, ns = non-significant).

### LAVR-289 is not toxic when administered once daily *per os* to immunosuppressed Syrian hamsters

In the accompanying manuscript we show that LAVR-289 is not toxic in mice when administered subcutaneously (see Marcheteau et al., same issue). To investigate if LAVR-289 is well tolerated *per os*, we used immunosuppressed Syrian hamsters that are an established model for adenovirus infection^24^. We tested vehicle alone (10% DMSO, 10% Cremophor, 80% PBS) or LAVR-289 at three dose levels of 25, 50 and 75 mg/kg, given orally once daily for 15 days. Groups (6 animals each) were followed daily for clinical signs and weight loss. No significant clinical observations were made in all groups. Hamsters in all groups gained weight at a similar rate (Fig. 3A). After 15 days of dosing, the animals were sacrificed and necropsied. No significant gross necropsy findings were noted for any of the groups. Also, plasma analysis on 26 different parameters did not show any impact of LAVR-289 on plasma composition at the end of the study (Table S1). These results show that LAVR-289 up to 75mg/kg is extremely well tolerated in immunosuppressed Syrian hamsters once daily using *per os* administration.

**Figure 3:**
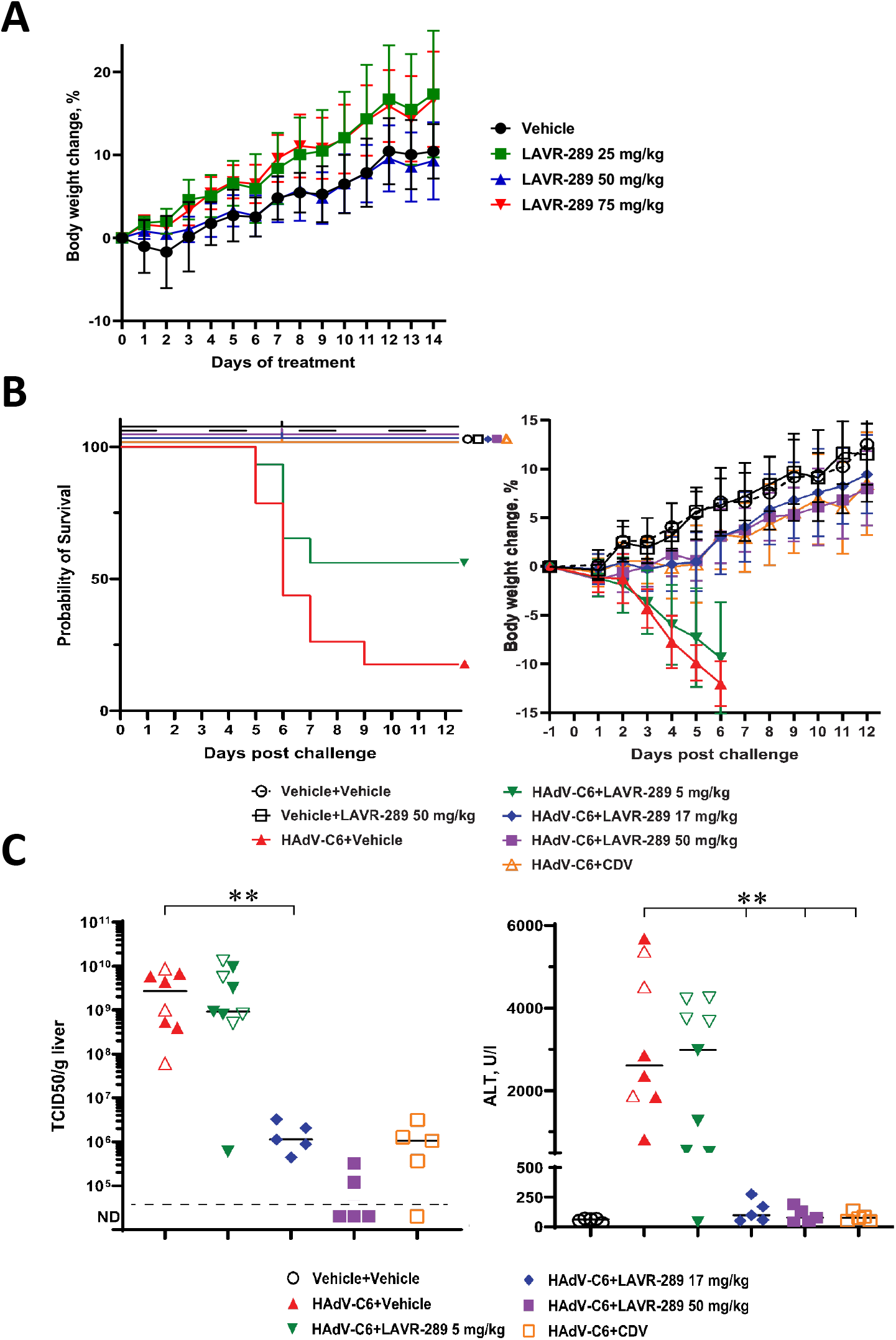
LAVR-289 given once daily by the oral route does not display any toxicity and protects hamster from HAdvC6 lethal challenge by reducing viral burden and liver damage. A) Weight curve of Syrian hamsters treated per os once daily for 14 days with LAVR-289 at the indicated concentrations. B) Survival probability curves (left) and weight curves (right) of hamsters infected intravenously with HAdv-C6 and treated with vehicle alone or LAVR-289 per os once daily at the indicated concentrations. CDV is used as a positive control. For weight curves, no group mean was calculated after an animal was sacrificed moribund from a given group. Virus burden assessed by TCID50 (C) and ALT levels (D) of the different groups. HAdV-C6+Vehicle vs. HAdV-C6+LAVR-289 5 mg/kg p=0.06, HAdV-C6+Vehicle vs. all other drug-treated p=0.0001 (Log-rank).

**Table S1.** Serum Chemistry.

| Group | ID | BUN | CREAT | AST | ALT | GGT | TBILI | ALP | AMY | LIP | GLU | CHOL | TRIG | CK | TP | ALB | GLOB | A/G | NA | K | Na/K | CA | CL | PHOS | ANION | OSMO | CO2 |
| --- | --- | --- | --- | --- | --- | --- | --- | --- | --- | --- | --- | --- | --- | --- | --- | --- | --- | --- | --- | --- | --- | --- | --- | --- | --- | --- | --- |
| Vehicle | 1 | 22.2 | 0.35 | 37.8 | 38.9 | 0.7 | 0.02 | 112 | 1559 | 36 | 173 | 105 | 149 | 198 | 6 | 3.4 | 2.2 | 1.51 | 141 | 10 | 14 | 15 | 97 | 10.6 | 0.1 | 307.4 | 54 |
|  | 2 | 19.9 | 0.3 | 49.3 | 37.2 | 0.8 | 0.02 | 90 | 1580 | 33 | 189 | 93 | 141 | 590 | 5 | 3.3 | 1.9 | 1.71 | 128 | 9.9 | 13 | 14 | 86 | 9.4 | 3.4 | 283.1 | 49 |
|  | 3 | 20.4 | 0.36 | 134 | 137 | 0.1 | 0.02 | 138 | 1679 | 162 | 151 | 106 | 164 | 754 | 6 | 3.4 | 2.2 | 1.57 | 139 | 8 | 17 | 14 | 92 | 9.3 | 3.4 | 298.1 | 52 |
|  | 4 | 21.1 | 0.31 | 32.1 | 44.1 | 0.4 | 0.02 | 89.4 | 1312 | 32 | 194 | 98 | 158 | 182 | 5 | 3.2 | 2.2 | 1.44 | 125 | 9.3 | 13 | 14 | 82 | 10.3 | 0.4 | 277.1 | 52 |
|  | 6 | 20.9 | 0.31 | 76.6 | 76.6 | 0.3 | 0.02 | 111 | 1551 | 34 | 155 | 93 | 134 | 703 | 5 | 3.1 | 2.1 | 1.52 | 126 | 9.4 | 13 | 13 | 85 | 10.4 | 3.9 | 276.9 | 47 |
| LAVR-289<br>25<br>mg/kg | 7 | 18.5 | 0.32 | 68.7 | 45.4 | 0.5 | 0.02 | 110 | 1281 | 30 | 193 | 100 | 164 | 543 | 5 | 3.1 | 1.7 | 1.79 | 119 | 9.6 | 12 | 13 | 79 | 8.9 | 0 | 265.5 | 50 |
|  | 8 | 19.7 | 0.3 | 91.2 | 157 | 0.3 | 0.01 | 101 | 908 | 42 | 175 | 100 | 165 | 271 | 5 | 3.1 | 2.2 | 1.45 | 124 | 9.2 | 13 | 13 | 84 | 9.4 | 1.5 | 273.5 | 48 |
|  | 9 | 20.1 | 0.34 | 50 | 63.3 | 0.1 | 0.02 | 104 | 1810 | 34 | 226 | 108 | 146 | 341 | 6 | 3.3 | 2.4 | 1.38 | 130 | 9.6 | 14 | 14 | 86 | 10.9 | 5 | 288.4 | 49 |
|  | 10 | 20.9 | 0.3 | 241 | 64.5 | 0.2 | 0.01 | 124 | 1689 | 37 | 100 | 134 | 160 | 1835 | 6 | 3.8 | 2.3 | 1.61 | 141 | 8.5 | 17 | 14 | 95 | 11.6 | 7.6 | 300.1 | 47 |
|  | 11 | 22 | 0.33 | 52.3 | 51.2 | 0.9 | 0.03 | 108 | 1314 | 38 | 215 | 114 | 148 | 404 | 6 | 3.3 | 2.2 | 1.55 | 126 | 9.2 | 14 | 14 | 83 | 10.1 | 6.2 | 280.3 | 46 |
|  | 12 | 21 | 0.32 | 197 | 189 | 0.4 | 0.01 | 121 | 1521 | 37 | 137 | 125 | 162 | 1296 | 6 | 3.6 | 2.3 | 1.57 | 141 | 8 | 18 | 13 | 94 | 10.8 | 9.5 | 301.3 | 46 |
| LAVR-289<br>50<br>mg/kg | 13 | 19 | 0.35 | 36.1 | 46.2 | 1 | 0.04 | 101 | 1542 | 33 | 219 | 110 | 169 | 239 | 6 | 3.5 | 2.2 | 1.56 | 142 | 10 | 14 | 15 | 97 | 10.9 | 3.5 | 311.4 | 52 |
|  | 14 | 19.4 | 0.33 | 111 | 49.3 | 0.3 | 0.02 | 115 | 1784 | 36 | 119 | 98 | 163 | 1019 | 6 | 3.4 | 2.5 | 1.4 | 141 | 8.3 | 17 | 14 | 92 | 10.8 | 10 | 300.2 | 47 |
|  | 15 | 25.9 | 0.38 | 58.5 | 46.5 | 1.7 | 0.01 | 127 | 1695 | 37 | 191 | 159 | 141 | 766 | 6 | 3.5 | 2.3 | 1.54 | 131 | 10 | 13 | 14 | 86 | 8.7 | 2.6 | 291.5 | 53 |
|  | 16 | 19.5 | 0.35 | 159 | 136 | 0.6 | 0.02 | 128 | 1510 | 37 | 147 | 120 | 153 | 809 | 6 | 3.4 | 2.2 | 1.55 | 137 | 9.2 | 15 | 14 | 90 | 10.2 | 8.8 | 296.1 | 47 |
|  | 17 | 16.8 | 0.32 | 39.8 | 42.9 | 1 | 0.01 | 106 | 1419 | 36 | 194 | 99 | 155 | 315 | 5 | 3.3 | 2 | 1.62 | 128 | 9.4 | 14 | 13 | 86 | 9.6 | 4.8 | 281.3 | 47 |
|  | 18 | 19.6 | 0.32 | 246 | 268 | 0.2 | 0.01 | 116 | 1631 | 37 | 233 | 124 | 169 | 2967 | 6 | 3.7 | 2.3 | 1.62 | 139 | 9 | 15 | 14 | 94 | 10.2 | 13.8 | 304.2 | 40 |
| LAVR-289<br>75<br>mg/kg | 19 | 21.7 | 0.36 | 40.3 | 48.6 | 0.5 | 0.02 | 99.4 | 1489 | 33 | 180 | 97 | 145 | 225 | 5 | 3.1 | 2.1 | 1.45 | 128 | 11 | 12 | 14 | 85 | 11.3 | 2.8 | 284.4 | 51 |
|  | 20 | 16.6 | 0.31 | 30.9 | 36.1 | 0.8 | 0.02 | 90.5 | 1474 | 31 | 162 | 89 | 156 | 153 | 5 | 3.1 | 2.2 | 1.38 | 122 | 9.1 | 13 | 13 | 81 | 9.6 | 4.5 | 267.8 | 46 |
|  | 21 | 20.2 | 0.34 | 37.1 | 51.7 | 0.6 | 0.01 | 94.3 | 1740 | 33 | 194 | 95 | 153 | 325 | 6 | 3.4 | 2.1 | 1.63 | 127 | 9.4 | 14 | 13 | 84 | 8.9 | 6.3 | 280.7 | 46 |
|  | 22 | 20.5 | 0.31 | 170 | 318 | 0.4 | 0.02 | 90.7 | 1542 | 33 | 179 | 91 | 144 | 573 | 5 | 3 | 1.8 | 1.65 | 127 | 9.3 | 14 | 13 | 85 | 9.5 | 3.6 | 279.8 | 48 |
|  | 23 | 19.4 | 0.32 | 39.2 | 44.4 | 0.8 | 0.01 | 103 | 1419 | 31 | 179 | 91 | 166 | 423 | 5 | 2.9 | 1.9 | 1.57 | 126 | 10 | 12 | 13 | 84 | 10.3 | 4.7 | 279 | 47 |
|  | 24 | 18 | 0.31 | 32.9 | 47.8 | 0.6 | 0.04 | 92.3 | 1333 | 36 | 220 | 116 | 173 | 179 | 5 | 3.1 | 2 | 1.59 | 121 | 10 | 12 | 13 | 80 | 9.2 | 8.9 | 271.5 | 42 |

### LAVR-289 *per os* administration once daily suppresses HAdV-C6 replication, liver damage and prevents animal death

To investigate if LAVR-289 is able to prevent or mitigate HAdV-induced pathology, we used immunosuppressed Syrian hamsters infected intravenously by HAdV-C6, a clinically relevant species C adenovirus. Under these conditions, HAdV-C6 replicates rapidly in the liver, triggering increased serum alanine aminotransferase (ALT) levels and animals die from fulminant hepatitis. The animals were distributed into 7 groups of 15 hamsters each, with the exception of LAVR-289 alone (without HAdV-C6 infection), which consisted of 9 animals. The hamsters were injected i.v. with vehicle or 2×10^10^ PFU/kg of HAdV-C6 and then treated with drug vehicle or LAVR-289 (p.o. q.d.). A control group received cidofovir (37 mg/kg IP, and then 20 mg/kg, 3 times weekly). For all groups, drug administration started one day before challenge. Infection of untreated hamsters induced mortality starting at 5 days post challenge (Fig 3B, left) and weight loss starting at 3 days post challenge (Fig 3B, right). Ten HAdV-C6-infected animals died in the vehicle conditions and 5 infected animals died at the 5 mg/kg LAVR-289. Necropsy indicated severe adenovirus pathology. Animals that survived in the 5mg/kg group recovered slowly, indicating that the dose of 5 mg/kg triggered partial protection in this challenge (Fig 3B, left). Doses of 17 mg/kg and 50 mg/kg prevented weight loss and all animals survived the challenge. No weight loss was observed with vehicle alone or animals treated with LAVR-289 at 50 mg/kg and left uninfected. To investigate if LAVR-289 in this experiment triggered a decrease in virus replication and liver damage, 5 animals of each group (except LAVR-289 50mg/kg uninfected) were sacrificed at 5 days post challenge, and the HAdV-C6 virus burden in the liver was examined by TCID50 assay. To assess liver damage, serum ALT levels were determined. As expected, in the untreated group, HAdV-C6 replication was pronounced, reaching 1.5×10^9^ TCID50 per g of liver (Fig 3C, left). The virus replication induced liver damage, with a high level of circulating ALT reaching around 2600 U/L, whereas uninfected controls are in the normal range (Fig 3C, right). Presence of LAVR-289 at 5mg/kg, despite having a trend in decreasing mortality, did not show significant reduction in virus burden or ALT levels (Fig 3C, left/right). LAVR-289 at 17mg/kg triggered a 99,99% decrease in virus levels in the liver and prevented liver damage. The same results were obtained for the CDV group. Some animals in the LAVR-289 at 50mg/kg group did not display any detectable infectious particles or elevated ALT levels.

### Discussion and summary

Adenovirus infection is usually self-limiting and benign in immunocompetent individuals but can cause serious disease in immunocompromised patients and during childhood. Even if a vaccine targeting two serotypes exists, it is limited to military applications. Nucleoside based antivirals in general are the most used inhibitors in the clinic because they are known to possess the highest barrier to resistance. Current nucleoside analogues used to treat HAdV infection are limited. Cidofovir and its pro-drug BCDV has been used to treat HAdV infection, but displayed high secondary symptoms and absence of activity in phase II clinical trial over placebo, respectively^10,25,26^. Ribavirin and Ganciclovir have also been used with limited efficacy. The broad variety of Adenovirus types use different entry receptors, target different tissues, and trigger different pathologies. Having a broad-spectrum antiviral inhibiting all types will thus be a great asset for treatment of HAdV-related pathologies. In this study, we have tested the activity of LAVR-289, a new acyclonucleoside phosphonate displaying antiviral activities on poxviruses and herpesviruses. We have shown that LAVR-289 inhibits the replication of HAdV-C5 ANCHOR virus at an EC50 of circa 100 nM and is as efficient as BCDV *in vitro*. Mechanistic studies have shown that LAVR-289 does not impact virus entry or genome release in the nucleus. In contrast, LAVR-289 prevents the formation of replication centers as shown by the absence of DBP clustering and the accumulation of replicated DNA indicated by the absence of ANCHOR marked replication centers. This likely indicates that LAVR-289 targets viral DNA replication, as we have shown for poxviruses. In turn, the absence of viral DNA replication prevents late viral gene expression, which is dependent on genome replication. Because late genes encode the structural proteins, virus propagation is inhibited. *In vivo* studies in immunosuppressed Syrian hamsters have shown that LAVR-289 is extremely well tolerated *in vivo* when administered once daily *per os*. The compound inhibited virus replication *in vivo*, protected the liver from adenovirus-induced pathology, and resulted in survival of 100% of treated animals even in the absence of a functional immune system. The broad-spectrum activity of LAVR-289 on different members of the dsDNA virus family can provide an effective tool for the treatment of immunocompromised people. For example, immunosuppression either induced (during grafting) or acquired (in HIV-1 positive individuals) can trigger the reactivation and/or infection with several viruses, such as herpesviruses and adenoviruses^27,28^. LAVR-289 administration in this case could impact simultaneously the replication of different virus families and offers the possibility to treat infection from different virus families at the same time with a single drug entity.

## Author Contributions

Conceptualization, K.T, H.W and F.G; Experiments, C.Q.F., A.T., A.C.S., S.K.G., N.L., V.R. L.A.A; Supervision, K.T., H.W and F.G. ;Writing—original draft,F.G. All authors have read and agreed to the published version of the manuscript.

## Acknowledgments

The authors thank Heather Greenstone (NIH NIAID - Division of Microbiology and Infectious Diseases) for continuous support.

